# Mesenchymal stem cells support human vascular endothelial cells to form vascular sprouts in human platelet lysate-based matrices

**DOI:** 10.1101/2022.01.31.478452

**Authors:** Eva Rossmanith, Sabrina Summer, Markus Pasztorek, Constantin Fiedler, Marion Gröger, Sabine Rauscher, Viktoria Weber, Michael B. Fischer

## Abstract

During tissue regeneration, mesenchymal stem cells can support endothelial cells in the process of new vessel formation. For a functional interaction of endothelial cells with mesenchymal stem cells a vascular inductive microenvironment is required. Using a cellular model for neo-vessel formation, we could show that newly formed vascular structures emanated from the embedded aggregates, consisting of mesenchymal stem cells co-cultured with human umbilical vein endothelial cells, into the avascular HPL-based matrices bridging distances up to 5 mm to join with adjacent aggregates with the same morphology forming an interconnected network. These newly formed vascular sprouts showed branch points and generated a lumen as sign of mature vascular development. Mesenchymal stem cells in fluid phase could bind to and interact with adherent human umbilical vein endothelial cells when a shear-force of 2 dyne/mm^2^ was applied in the flow chamber using a Bioflux®200 device. Under these conditions, mesenchymal stem cells bind to human umbilical vein endothelial cells previously damaged by laser beam irradiation to interact and shed particles.

In conclusion, we observed that mesenchymal stem cells support human umbilical vein endothelial cells to form new vessels in HPL-based matrices when co-cultured in spherical aggregates to generate complex vascular networks in a primarily avascular scaffold and bind to the damaged cells under shear-force to eventually aid in their regeneration.

## Introduction

Mesenchymal stem cells (MSCs) are the stem cells of the connective tissue showing strong regenerative activity to counteract degenerative diseases due to their multi-lineage differentiation and self-renewal capability [1–4]. MSCs support connective tissue regeneration by integration into the damaged site and their maturation along the mesenchymal lineage towards organ specific cells. MCS can also release their secretome or shed microvesicles (MVs) and exosomes into the damaged tissue or donate mitochondria to rescue damaged cells [5, 6]. Finally, MSCs contribute to tissue and organ regeneration by supporting endothelial cells (ECs) in the process of new vessel formation [7–12]. Sprouting angiogenesis in response to ischemia and hypoxia starts with extracellular matrix (EC) degradation and detachment of perivascular MSCs from capillaries in the affected area. This allows the endothelial tip cell to become invasive and induce formation of filopodia and lamellipodia in response to guidance cues determining the direction of vascular growth [13–15]. After initiation, stalk cells that lie behind the tip cells tend to proliferate and extend the vessel by forming ECM, junctions, and lumen [14, 15]. Once the tip cells anastomose with other tip cells to form a potential circuit, blood vessel maturation takes place, which involves the recruitment of endothelial precursors from the circulation [14]. This favors deposition of ECM by ECs and perivascular MSCs and the initiation of a blood flow [13]. Under these conditions perivascular MSCs come into direct contact with immature capillaries to stabilize the newly formed tubular network by abluminal coverage [15]. Vascular basement membranes can directly interact with ECs that line the inside of the newly formed blood vessel and perivascular MSCs covering the outside [15]. MSCs can produce and deposit collagen type IV, a major constituent of vascular basement membranes, within newly formed blood vessels both *in vivo* and in fibrin-based matrices [16, 17]. During the integration into the vascular wall MSCs respond to EC-derived TGF-β [18, 19]. Contact-dependent activation of TGF-β is a common driver of MSC differentiation into vascular smooth muscle cells during vessel maturation, involving TGF-β/ALK signaling [19].

Important for an MSC-mediated support of ECs in angiogenesis models is the tissue source MSCs are isolated from, their cultivation conditions and the scaffold composition [1–4]. Among different EC types investigated, HUVECs were extensively studied in co-cultures with MSCs, smooth muscle cells, or pericytes as supportive cells [7–12]. While bone marrow-derived and adipose-derived MSCs supported HUVEC organization into vascular-like structures, amniotic fluid-derived MSCs appeared to provide even better pro-vascular support, indicating the importance of an earlier stage in ontogeny [8, 12]. Cultivation of MSCs in adherence to plastic surface combined with xeno-based medium supplements could substantially modify the phenotype and transcriptional activity, resulting in an impaired interaction with HUVECs in angiogenic models [20]. In search of xeno-free cultivation solutions, HPL-based medium supplements were used to maintain the phenotype of cultured MSCs, reduce stress fiber formation and preserve mitochondrial function [21]. However, current information addressing the role of HPL-based matrices on the communication between ECs and MSCs during new-vessel formation is limited.

Here, we investigated the potential of amnion-derived MSCs to support HUVECs in the process of new-vessel formation using primary avascular HPL-based matrices in an autologous setting. Those HPL-based scaffolds combine fibrin-based matrices with the bioactive content of platelets [22–24]. Spherical aggregates containing MSCs and HUVECs, generated by the hanging drop technology, were embedded into 20% HPL to investigate new vessel formation. As developmental signs of vascular maturation, the capability of these newly formed vascular structures to generate tube-like structures and bridge distances of up to 5 mm to the neighbor spherical aggregates with the same morphology was observed. Finally, using a commercially available microfluidic system, the potential of MSCs to bind to and interact with a layer of adherent HUVECs was examined under flow conditions applying the Bioflux®200 device.

## Materials and Methods

### Isolation, cultivation, and characterization of MSCs and HUVECs

Placental tissues were obtained from healthy delivering women in accordance with the Austrian Hospital Act (KAG 1982) and studies were ethically confirmed (GSl-EK-4/3122015). Amnion-derived MSCs from placental tissue were isolated and characterized with CD73-APC, CD90-FITC and CD105-PE-Cy7 (all from eBioscience, San Diego, CA) by flow cytometry (CytoFLEX XL, Beckman Coulter GmbH, Krefeld, Germany) [21] and HUVECs were isolated from the umbilical cord and characterized as published previously [25].

### Generation of spherical aggregates in hanging drops

Spherical aggregates were generated by co-cultivation of 4500 MSCs and 500 HUVECs (isolated from the same placenta) on lids of petri dishes (Greiner Bio-One, Kremsmünster, Austria) using the hanging drops technology [21]. Cells were co-cultivated in a volume of 25 μl M-199 medium supplemented with 10% FBS (Gibco, Thermo Fisher Scientific), endothelial growth supplement (20 μg/ml, Becton Dickinson, Franklin Lakes, NJ) and Heparin (10 IU/ml, Baxalta, Vienna, Austria). Alternatively, MSC aggregates of 5000 MSCs were cultured in MSC-BM^TM^ or MSC-GM^TM^ (both from Lonza Group Ltd, Basel, Switzerland) using the same technology and monitoring of aggregate formation was performed by phase contrast microscopy (IMT2, Olympus Austria GmbH, Vienna) equipped with a digital camera (DP50, Olympus).

### Imaging of spherical aggregates by scanning electron microscopy

MSC aggregates were adhered on Nunc Thermanox^TM^ coverslips (Nunc, Thermo Fisher Scientific), fixed, dehydrated, mounted on conductive double side adhesive carbon tabs (Miere to Nano V.O.F., Haarlem, Netherlands), sputtered with gold and analyzed by a scanning electron microscope (FlexSEM, Hitachi Ltd. Corp., Tokyo, Japan) as previously described [21].

### Imaging of spherical aggregates by confocal microscopy

MSC aggregates were cultivated in Nunc^TM^ Lab-Tek^TM^ II chamber slides (Nunc, Thermo Fisher Scientific), fixed and permeabilized after 16 h and stained with Alexa Fluor AF® 594 phalloidin (0.1 U/ml, Molecular Probes, Thermo Fisher Scientific) to reveal filamentous (f)-actin. To stain focal adhesions a mouse mAb to paxillin (2 μg clone B2, Santa Cruz Biotechnology, Dallas, TX) followed by goat-anti-mouse Fab fragments labeled with AF® 488 (3 μg/ml, Jackson Laboratories, Bar Harbor, MN) were used and nuclei were stained with DAPI (Sigma-Aldrich, St. Louis, MI). The slides were mounted with Fluoromount-G^TM^ (Southern Biotechnology, Thermo Fisher Scientific) and analyzed with an Apochromat 63x objective on a confocal microscope (TCS SP8, Leica Microsystem GmbH, Wetzlar, Germany) using the LAS X-software [21].

### Extracellular matrix composition of spherical aggregates by confocal microscopy

MSC aggregates were embedded in OCT compound (Tissue Tek, Sakura Finetek Europe BV, Alphen an den Rijn, Netherlands) and shock frozen in liquid nitrogen to generate cryosections of 4 μm on a Cryostar NX70 (Thermo Fisher Scientific). Sections were mounted on Superfrost Plus slides (Menzel Glas, VWR, Thermo Fisher Scientific), fixed in 4 °C acetone and stored at −80 °C. ECM components were detected using mouse mAbs specific for collagen type I (0.5 μg/ml clone ab90395, Abcam®, Cambridge, UK), collagen type IV (2 μg /ml clone M0785, Dako, Agilent Pathology Solutions, Santa Clara, CA), fibronectin (2 μg/ml clone MAB1918, R&D Systems), or a rabbit mAb specific for laminin (5 μg/ml clone ab11575, abcam®, Cambridge, UK). Primary mAbs were subsequently stained either with goat anti-mouse lgG AF® 594 (1:500) or with goat anti-rabbit lgG AF® 594 (1:500, Jackson Laboratories, US). MSCs were counter-stained with CD90-FITC and nuclei were stained with DAPI. The slides were mounted with Fluoromount-G^TM^ (Southern Biotechnology, US) and analyzed on a confocal microscope (SP8, Leica, Germany) [21].

### Vascular network formation in HPL-based matrices by confocal microscopy

Human thrombin (20 U/ml, Sigma-Aldrich, Germany) was added to 20% HPL (MacoPharma, France) in M-199 Medium supplemented with endothelial cell growth supplement (ECGS) and 500 μl of the mix was pipetted onto an ibidi μ-Dish (ibidi GmbH, Germany). After gel formation occurred, spherical aggregates consisting of MSCs and HUVECs were embedded on the gel surface at positions with 2-5 mm between the aggregates (Fig 3 scale bar). After 48 h 500 μl of M-199 medium supplemented with 8% HPL and ECGS was added that was exchanged once a week. New vessel formation was monitored twice a week using the ChemiDoc system (Bio-Rad Laboratories Inc., US) and phase contrast microscopy (IMT-2, Olympus). After 21 d of cultivation, HPL-based gels were fixed with 4% formaldehyde overnight to guarantee gel integrity. Cells within the HPL-based gels were than stained with AF® 488 phalloidin (0.1 U/ml, Molecular Probes, US) and HUVECs with a rabbit anti-human von Willebrand factor (vWF) mAb (2 μg/ml, Dianova, German) followed by goat anti-rabbit lgG Alexa Fluor® 594 (1:500, Jackson Laboratories, US) to reveal the primary Ab. Nuclei were stained with DAPI. Alternatively, before spheroid formation by hanging drops technique, MSCs were labeled with Cell Tracker™ Green CMFDA dye (1:1000 dilution) and HUVECs were labeled with Cell Tracker™ Red CMTPX dye (1:1000 dilution, both from Thermo Fisher Scientific, US). The gels were covered with 500 μl PBS and z-stack analysis was performed using an Apochromat 10x objective and confocal microscopy (SP8, Leica, Germany).

### Binding of MSCs to adherent HUVECs in flow cells of the BioFlux®200 device

An electro pneumatically controlled BioFlux®200 system (Fluxion Biosciences, lnc., US) with microfluidic plates of 24 independent flow chambers with a dimension of 350 μm width, 1500 μm length and 70 μm height was used to investigate binding of MSCs to a HUVEC layer applying 2 dyne/cm^2^ flow. Flow channels were coated with fibronectin (2.5 μg/cm^2^, Gibco, Thermo Fisher Scientific) and 2*10^5^ HUVECs in a volume of 200 μl EGM-2 medium (Lonza Group Ltd, Switzerland) were seeded and cultivated overnight under static conditions. To reach confluency of the HUVEC layer, perfusion with 2 dyne/cm^2^ was applied for 24 to 48 h. Damage to the HUVEC layer in an area of 150 μm in diameter (0.0176 mm^2^) was introduced by a laser capture micro-dissector (LMD6, Leica Microsystems GmbH, Germany). After dead HUVECs were removed, MSCs were applied via the inlet terminal in a density of 1*10^5^ cells/ml under 2 dyne/mm^2^ flow rate. Binding of MSCs to the HUVEC layer was monitored by phase contrast microscopy and the reaction was stopped after 3 h by fixation with 4% formaldehyde. Flow cells were washed and incubated with mouse anti-human VE-cadherin (2 μg/ml clone 123413, R&D Systems, US) followed by goat anti-mouse lgG AF® 594 (1:500, Jackson Laboratories, US) to stain tight junctions of ECs and CD90 FITC (2.5 μg/ml, eBioscience, US) to stain MSCs. Nuclei were stained with DAPI. Binding of MSCs to the EC layer was investigated using an LSM 700LS confocal laser scanning microscope (Carl Zeiss Microscopy GmbH, Germany) and the ZEN software program (Zeiss, Germany). A reference area of 0.1 mm^2^ (elliptic area of 330 x 415.5 μm) was investigated by phase contrast microscopy to quantify MSC binding to the HUVEC layer.

## Results

### MSCs generate delicate actin filaments within 3D spherical aggregates and contribute to extracellular matrix for stability

MSCs of passage 1, expressing the mesenchymal specific markers CD73, CD90 and CD105, were used to generate spherical aggregates applying the hanging drops technology (Fig 1A and S1 Fig). We showed that 5000 MSCs formed stable 3D spherical aggregates using scanning electron microscopy and confocal microscopy (Fig 1B, C). When spherical MSC aggregates were adhered to plastic surface for imaging, MSCs emanated from the aggregates and adhered as single cells (Fig 1D). Those single MSCs developed ventral stress fibers that were anchored at both sides to focal adhesions, spanning the entire cell and showed actin branching as described previously (Fig 1E) [21].

**Fig 1.**
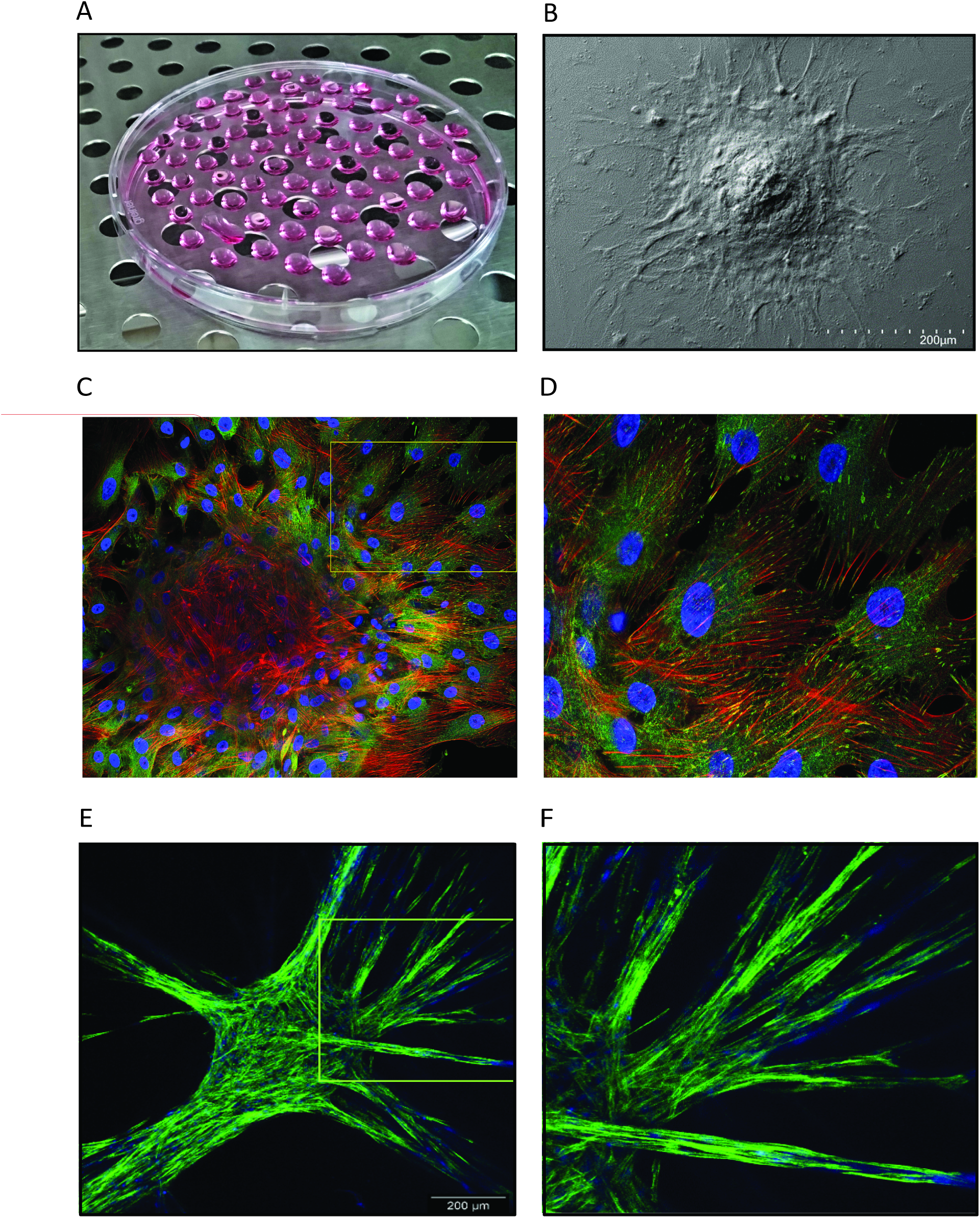
Spherical aggregate of MSCs. Spherical aggregates generated by cultivation of 5000 MSCs on the lid of a petri dish using the hanging drop technology (A). Representative image of a spherical MSC aggregate adherent to a coverslip obtained by scanning electron microscopy (B) and image of a spherical MSC aggregate stained with a mAb specific for paxillin to reveal focal adhesions (green), with phalloidin to stain actin filaments (red), and DAPI to highlight nuclei (blue) generated by laser scanning microscopy (C, D). Image of a spherical MSC aggregate embedded in HPL matrices showing sprout formation after 14 days of cultivation with MSC actin filaments stained with phalloidin (green) and nuclei with DAPI (blue) (E, F).

When spherical MSC aggregates were cultured in xeno-based medium, such as MSC-GM, MSCs generated collagen type I, collagen type IV, laminin, and fibronectin as major ECM components for aggregate stability (S2 Fig). The amount of individual ECM components produced by MSCs within the 3D spherical aggregates, however, did not change substantially during the period of 20 days of cultivation. When HPL was used as xeno-free medium supplement to culture MSC aggregates, the production of collagen type I, collagen type IV, laminin, and fibronectin by MSCs did not differ from ECM production of MSCs cultivated in xeno-based medium (S3 Fig).

### Vascular sprout formation in HPL-based matrices induced by spherical MSC/HUVEC aggregates

To study the supportive role of MSCs in the *de novo* vessel formation of HUVECs, we used 20% HPL-based scaffolds. Before 3D spherical aggregate formation, MSCs were labeled with CellTracker™ Green and HUVECs with CellTracker™ Red. The aggregates were embedded in HPL-based matrices in distances of 2-5 mm. A digital documentation system was used to document vascular development over a period of 20 days (Fig 2).

**Fig 2.**
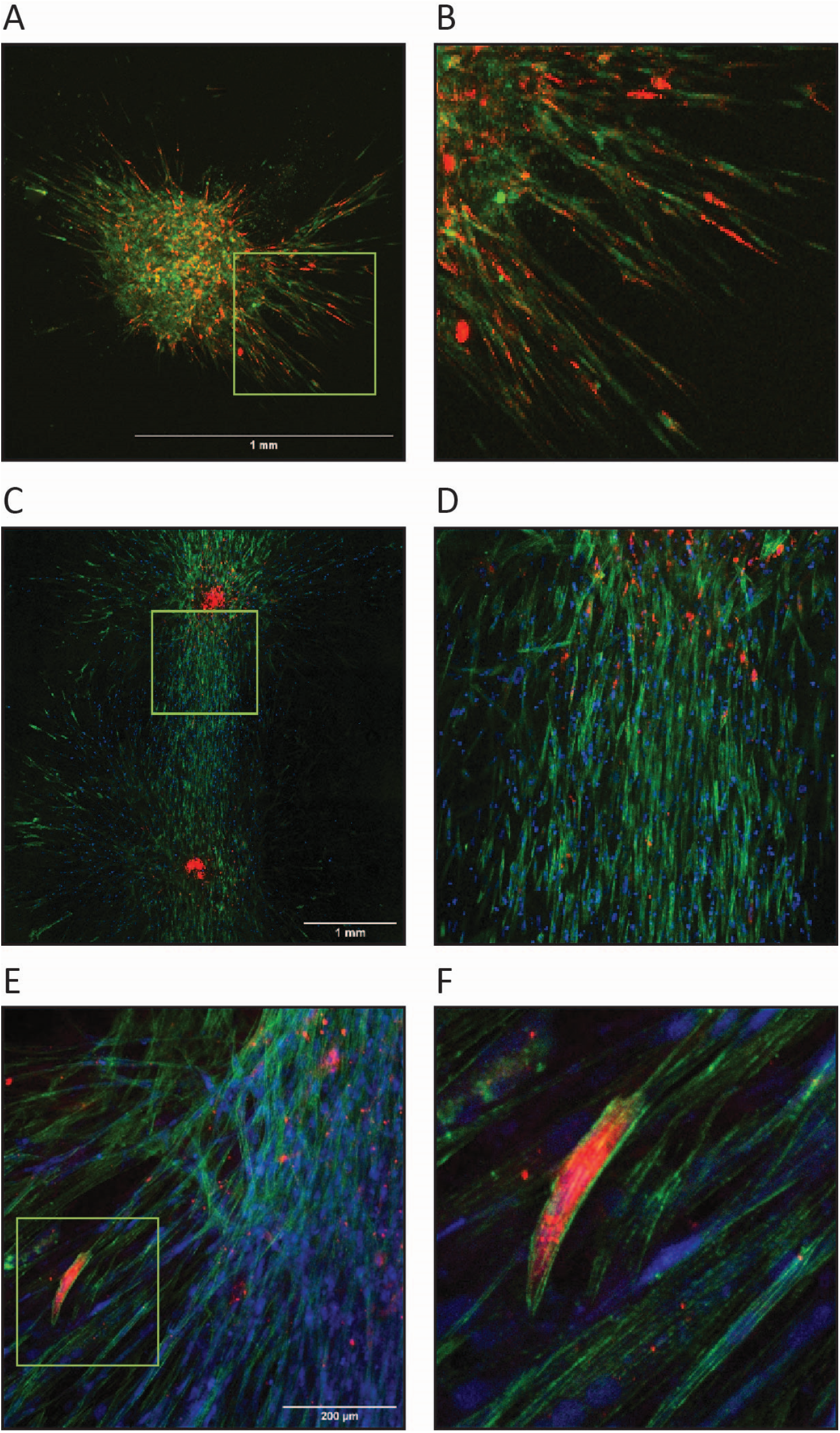
Spherical aggregates in HPL-based matrices. Spherical aggregates consisting of 4500 MSCs and 500 HUVECs were generated by the hanging drop technology and subsequently embedded in 20% HPL matrices. MSCs stained with CellTracker™ Green and HUVECs stained with CellTracker™ Red were co-cultured to generate spherical aggregates and these aggregates were embedded into HPL-based matrices for 6 days to see vascular sprouts emanating from the aggregate (A, B). A representative image of two aggregates embedded into the HPL-based matrices in a distance of approximately 3 mm showed interconnection of the sprouts after 21 days of cultivation (C, D). MSCs and HUVECs actin filaments were stained with phalloidin (green) and to segregate the two cell populations HUVECs we stained by the endothelial specific marker vWF (red) (C-F).

Here, we showed that the spatial organization of cellular outgrowths into the initially avascular translucent HPL-based matrices occurred within the first five days of cultivation (Fig 2A, B). These outgrowths changed their appearance to a spindle-shape cell morphology, forming cellular strands that continued to grow towards neighboring strands with the same morphology (Fig 2C, D). These newly formed sprouts developed in different planes within the HPL-based matrices to form cord-like structures with a mix of labeled HUVECs and MSCs (Fig 2E, F). Within these newly formed sprouts HUVECs showed nuclear elongation and occasionally formed tip cells positive for vWF (Fig 2E, F). Generation of vascular branch points and a lumen are important maturation steps of newly formed blood vessels during their establishment (Fig 2F).

Here, we showed that these newly formed vascular structures showed branch points and generated a lumen in our static system in the total absence of a blood circulation introducing flow (Fig 3E).

**Fig 3.**
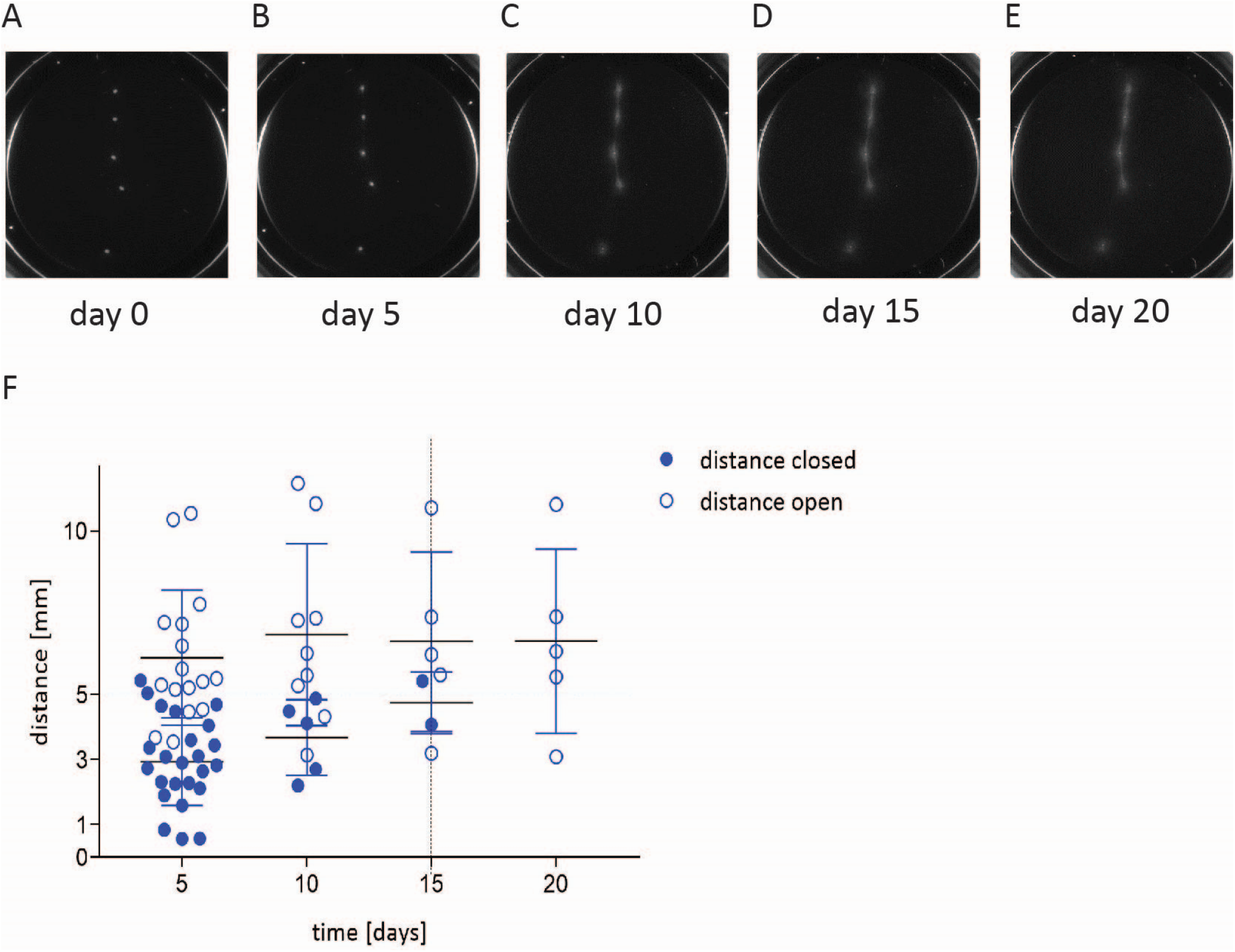
Vascular sprout formation in HPL-based matrices. The capability of two spherical aggregates consisting of 4500 MSCs and 500 HUVECs to bridge distances of up to 5 mm with their vascular sprouts was investigated using phase contrast microscopy (A-E). The closure time of two adjacent spherical aggregates was recorded over a period of 20 days. The joining of vascular sprouts from two adjacent spherical aggregates was recorded at the day indicated with successful joining (filled circles) and failure to close (open circles) (F).

No evidence was found for a mixed endothelial/mesenchymal phenotype or cross-differentiation of MSCs to endothelial cells in our experimental setting. The vascular sprouts that emanated from a spherical aggregate consisting of co-cultured MSCs and HUVECs could bridge distances up to 5 mm to form an interconnected network with adjacent aggregates showing the same morphology after 10 to 15 days of cultivation (Fig 3).

### Binding of MSCs to adherent HUVEC layer under flow

The capability of MSCs to bind to and interact with adherent HUVECs under dynamic fluid conditions in a commercially available microfluidic system was investigated using the BioFlux®200 device that enabled multiple temperature-controlled flow assays to run in parallel. MSCs applied in the fluid phase bound to the layer of adherent HUVECs in a density of 35-40 MSCs/0.1 mm^2^ when a flow rate of 2 dyne/cm^2^ was applied (Fig 4A). In case of cell damage induced by laser radiation, MSCs bound to the HUVECs within this area with a higher frequency (Fig 4B, D). MSCs could either adhere to the fibronectin-coated surface to communicate with HUVECs from a certain distance by long nano-tubular extrusions (Fig 4B) or directly to the HUVECs. Bound MSCs were able to shed particles into their immediate surrounding that could bind to the surface of HUVECs and certain bound MSCs showed active transmigration through the EC layer (Fig 4C).

**Fig 4.**
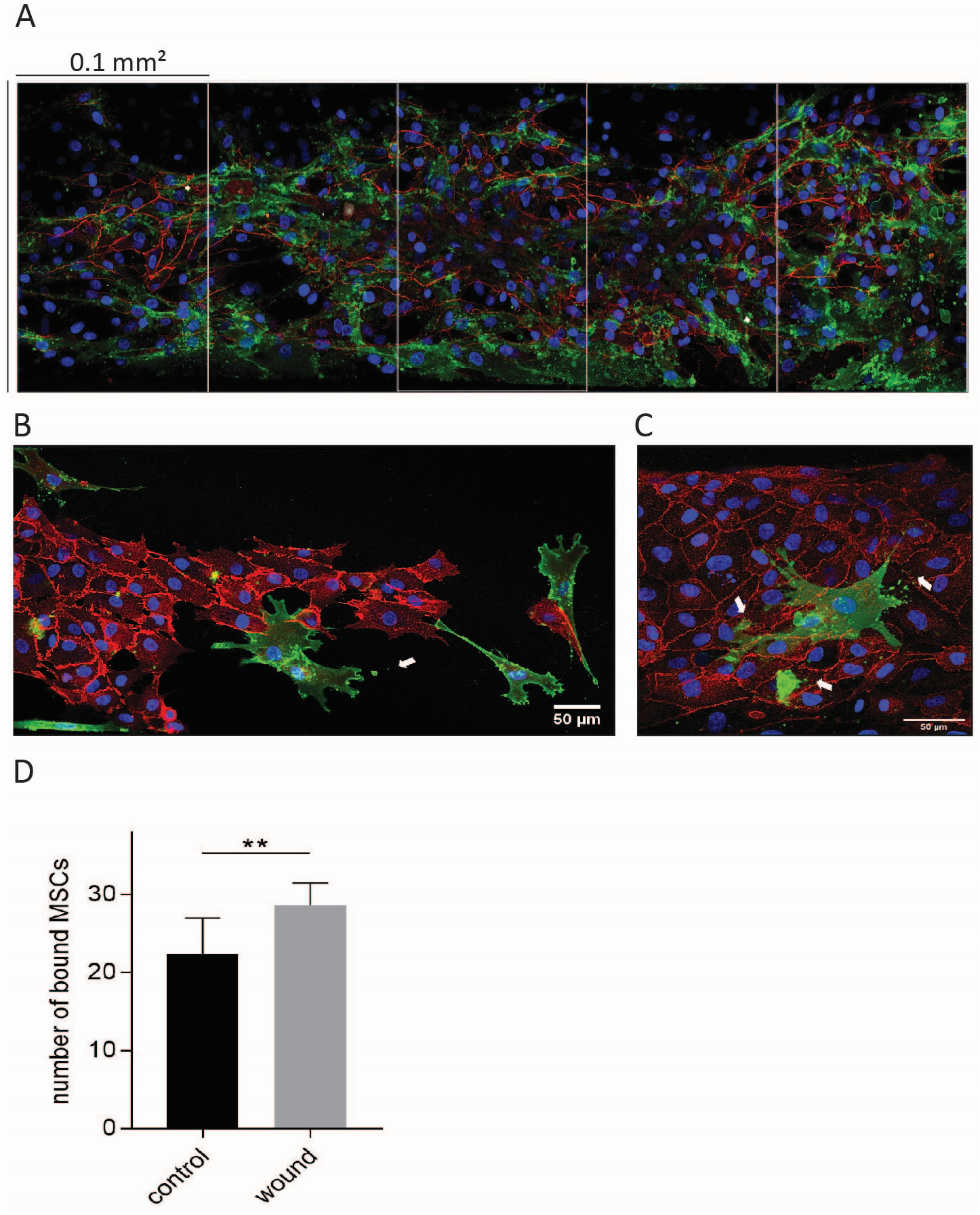
MSCs binding to a layer of HUVECs under flow in a microfluidic system. MSCs in fluid phase were applied under 2 dyne/mm2 shear rate to bind to a layer of HUVECs in a BioFlux®200 system with a total chamber area is 0.525 mm2 (1500 mm x 350 mm) and one sector indicating 0.1 mm2 (A). A representative area of damaged HUVECs stained with CD144 (VE-cadherin, red) and adherent MSCs stained with CD90 (green) (B). Bound MSCs (green) can shed particles (arrows) and can actively transmigrate under the HUVEC-layer (red) (C).

## Discussion

Tissue regeneration relies on neo-vessel formation, a process that requires MSCs to support ECs during initiation and development of vascular sprouts. Tissue damage with vascular injury leads to leakage of plasma components into the damaged site and the generation of an early provisional matrix consisting of fibrin-rich polymers with interspersed cross-linked fibronectin and platelets [26–29]. The clotting response and the release of platelet α-granule content into the fibrin-rich provisional matrix stop bleeding and create a basic scaffold for MSC and EC interaction [26]. Next to fibrinogen, more than 300 bioactive substances are released by activated platelets that can potentially interfere with MSC-EC communication [30]. We argued that fibrin-based matrices that incorporate platelet secretome serve as ideal scaffolds to create new vessels close to physiological conditions [26–29]. Here, we established an HPL-based matrix model system to investigate new vessel formation induced by HUVECs supported by MSCs. We found that spherical aggregates consisting of HUVEC co-cultured with MSCs served as ideal tools for vascular engineering since MSCs provided the ECM components necessary for aggregate stability, such as collagen type I, fibronectin, laminin, and collagen type IV [16, 17, 31]. This is of importance because the early provisional fibrin-rich matrix is replaced during the process of healing by locally produced fibronectin, laminin and proteoglycans generated by invading cells of the mesenchymal lineage [26, 32]. We demonstrated that MSCs within spherical aggregates provide collagen type I and IV as well as fibronectin and laminin important for aggregate stability, while HUVECs did not add to ECM formation under these conditions. The fibronectin assembly is driven by a complex process of cell binding, molecular extension of the protein through physical forces to expose multiple cryptic self-association sites [27, 28]. In the polymerized state, fibronectin can be considered as a scaffold signaling protein favoring new vessel formation [28, 29, 33–37]. In this study we showed the formation of vascular sprouts originating from the inserted spherical MSC/HUVEC aggregates into the primarily avascular HPL-based matrices. When spherical MSC/HUVEC aggregates were embedded into HPL gel at distances of up to 5 mm between the aggregates the sprouts grew towards the adjacent aggregate showing the same morphology and joined to a connected network. Interestingly, these outgrowths also formed a lumen, similar to results shown by Ruger et al [22].

When the capability of MSCs to specifically bind to HUVECs under dynamic conditions was investigated using the BioFlux®200 system, we found that MSCs could bind to adherent HUVECs under controlled shear flow of 2 dyne/cm^2^ [38]. The shear rates represent the physiological values in human venules with diameters ranging from 20–70 μm [39]. When the perfused HUVEC layer was damaged by UV laser beam irradiation, MSCs bound preferentially to the HUVECs located at the rim of damage. MSCs not only bound to HUVECs, but also generated long nano-tubular extrusions to contact with the damaged HUVECs from a certain distance without immediate binding. MSCs shed particles into their immediate surrounding that could bind to the cell surface of HUVECs, giving further evidence for active interaction (Fig 4B, C).

## Conclusion

Here, we provide evidence that spherical MSC/HUVEC aggregates can initiate new vessel formation and form vascular sprouts by self-induction in HPL-based matrices. These newly formed sprouts can bridge distances of up to 5 mm to join and form an interconnected network with an adjacent aggregate having the same morphology. Although we found no evidence for trans-differentiation of MSCs to ECs expressing CD31 or vWF in our system [35], we could show that MSCs support ECs in the initiation process of new vessel formation *in vitro.* Furthermore, MSCs can adhere to a layer of adherent HUVECs when a shear flow of 2 dyne/cm^2^ was applied to interact, shed extracellular vesicles, and eventually transmigrate. Our results give evidence for the capability of MSCs to interact actively with HUVECs either in the process of new vessel formation in artificial HPL-based matrices without addition of pro-vascular bioactive molecules or in a microfluidic system, applying shear flow. From the clinical view, these assays can give further information on the ability of MSCs to contribute to the regeneration of vascular defects.

## Acknowledgement

The authors thank the Austrian Cluster for Tissue Regeneration (ACTR) headed by Prof. DI Dr. Heinz Redl for networking. This work was financially supported by the Government of Lower Austria (WST3-F-5030664/009-2018) and co-funded by the European Regional development Fund (INTERREG AT-CZ 133).

## Supporting information

**S1 Fig. Phenotypic characterization of MSCs.** MSCs were phenotypically characterized by flow cytometry using the stem cell markers CD73, CD90 and CD105.

**S2 Fig. Extracellular matrix formation by MSCs in spherical aggregate cultivated in MSC-GM.** Spherical MSC aggregates were cultivated for the time indicated (3d, 6d, 9d and 12d) and cryo-sectioned and stained with mAbs specific form collagen type I (A-C), collagen type IV (D-G), laminin (H-K) and fibronectin (I-L) (red) and counterstained with CD90 (green). For nuclei staining DAPI was applied (blue).

**S3 Fig. Extracellular matrix formation by MSCs in spherical aggregate cultivated in MSC-BM with 8% HPL.** Spherical MSC aggregates were cultivated for the time indicated (3d, 6d, 9d and 12d), cryo-sectioned and stained with mAbs specific form collagen type I (A-C), collagen type IV (D-G), laminin (H-K) and fibronectin (I-L) (red) and counterstained with CD90 (green). For nuclei staining DAPI was applied (blue).

## References

1. Sagaradze GD, Basalova NA, Efimenko AY, et al. Mesenchymal Stromal Cells as Critical Contributors to Tissue Regeneration. Front Cell Dev Biol 2020; 8: 576176. 2020/10/27. DOI: 10.3389/fcell.2020.576176.

2. Shammaa R, El-Kadiry AE, Abusarah J, et al. Mesenchymal Stem Cells Beyond Regenerative Medicine. Front Cell Dev Biol 2020; 8: 72. 2020/03/07. DOI: 10.3389/fcell.2020.00072.

3. Wu X, Jiang J, Gu Z, et al. Mesenchymal stromal cell therapies: immunomodulatory properties and clinical progress. Stem Cell Res Ther 2020; 11: 345. 2020/08/11. DOI: 10.1186/s13287-020-01855-9.

4. Bianco P. Back to the future: moving beyond “mesenchymal stem cells”. J Cell Biochem 2011; 112: 1713–1721. 2011/03/19. DOI: 10.1002/jcb.23103.

5. Baglio SR, Pegtel DM and Baldini N. Mesenchymal stem cell secreted vesicles provide novel opportunities in (stem) cell-free therapy. Front Physiol 2012; 3: 359. 2012/09/14. DOI: 10.3389/fphys.2012.00359.

6. Li C, Cheung MKH, Han S, et al. Mesenchymal stem cells and their mitochondrial transfer: a double-edged sword. Biosci Rep 2019; 39 2019/04/14. DOI: 10.1042/BSR20182417.

7. Merfeld-Clauss S, Gollahalli N, March KL, et al. Adipose tissue progenitor cells directly interact with endothelial cells to induce vascular network formation. Tissue Eng Part A 2010; 16: 2953–2966. 2010/05/22. DOI: 10.1089/ten.TEA.2009.0635.

8. Roubelakis MG, Tsaknakis G, Pappa KI, et al. Spindle shaped human mesenchymal stem/stromal cells from amniotic fluid promote neovascularization. PLoS One 2013; 8: e54747. 2013/01/30. DOI: 10.1371/journal.pone.0054747.

9. Lin RZ, Moreno-Luna R, Zhou B, et al. Equal modulation of endothelial cell function by four distinct tissue-specific mesenchymal stem cells. Angiogenesis 2012; 15: 443–455. 2012/04/25. DOI: 10.1007/s10456-012-9272-2.

10. Au P, Tam J, Fukumura D, et al. Bone marrow-derived mesenchymal stem cells facilitate engineering of long-lasting functional vasculature. Blood 2008; 111: 4551–4558. 2008/02/08. DOI: 10.1182/blood-2007-10-118273.

11. Traktuev DO, Prater DN, Merfeld-Clauss S, et al. Robust functional vascular network formation in vivo by cooperation of adipose progenitor and endothelial cells. Circ Res 2009; 104: 1410–1420. 2009/05/16. DOI: 10.1161/CIRCRESAHA.108.190926.

12. Verseijden F, Posthumus-van Sluijs SJ, Pavljasevic P, et al. Adult human bone marrow- and adipose tissue-derived stromal cells support the formation of prevascular-like structures from endothelial cells in vitro. Tissue Eng Part A 2010; 16: 101–114. 2009/08/01. DOI: 10.1089/ten.TEA.2009.0106.

13. Ribatti D and Crivellato E. “Sprouting angiogenesis”, a reappraisal. Dev Biol 2012; 372: 157–165. 2012/10/04. DOI: 10.1016/j.ydbio.2012.09.018.

14. Siekmann AF, Affolter M and Belting HG. The tip cell concept 10 years after: new players tune in for a common theme. Exp Cell Res 2013; 319: 1255–1263. 2013/02/20. DOI: 10.1016/j.yexcr.2013.01.019.

15. Makanya AN, Hlushchuk R and Djonov VG. Intussusceptive angiogenesis and its role in vascular morphogenesis, patterning, and remodeling. Angiogenesis 2009; 12: 113–123. 2009/02/06. DOI: 10.1007/s10456-009-9129-5.

16. Somaiah C, Kumar A, Mawrie D, et al. Collagen Promotes Higher Adhesion, Survival and Proliferation of Mesenchymal Stem Cells. PLoS One 2015; 10: e0145068. 2015/12/15. DOI: 10.1371/journal.pone.0145068.

17. Yang C, DelRio FW, Ma H, et al. Spatially patterned matrix elasticity directs stem cell fate. Proc Natl Acad Sci U S A 2016; 113: E4439–4445. 2016/07/21. DOI: 10.1073/pnas.1609731113.

18. Carvalho RL, Jonker L, Goumans MJ, et al. Defective paracrine signalling by TGFbeta in yolk sac vasculature of endoglin mutant mice: a paradigm for hereditary haemorrhagic telangiectasia. Development 2004; 131: 6237–6247. 2004/11/19. DOI: 10.1242/dev.01529.

19. Hirschi KK, Rohovsky SA and D’Amore PA. PDGF, TGF-beta, and heterotypic cell-cell interactions mediate endothelial cell-induced recruitment of 10T1/2 cells and their differentiation to a smooth muscle fate. J Cell Biol 1998; 141: 805–814. 1998/06/13. DOI: 10.1083/jcb.141.3.805.

20. Kocaoemer A, Kern S, Kluter H, et al. Human AB serum and thrombin-activated platelet-rich plasma are suitable alternatives to fetal calf serum for the expansion of mesenchymal stem cells from adipose tissue. Stem Cells 2007; 25: 1270–1278. 2007/01/27. DOI: 10.1634/stemcells.2006-0627.

21. Pasztorek M, Rossmanith E, Mayr C, et al. Influence of Platelet Lysate on 2D and 3D Amniotic Mesenchymal Stem Cell Cultures. Front Bioeng Biotechnol 2019; 7: 338. 2019/12/06. DOI: 10.3389/fbioe.2019.00338.

22. Ruger BM, Buchacher T, Giurea A, et al. Vascular Morphogenesis in the Context of Inflammation: Self-Organization in a Fibrin-Based 3D Culture System. Front Physiol 2018; 9: 679. 2018/06/21. DOI: 10.3389/fphys.2018.00679.

23. Ruger BM, Buchacher T, Dauber EM, et al. De novo Vessel Formation Through Cross-Talk of Blood-Derived Cells and Mesenchymal Stromal Cells in the Absence of Pre-existing Vascular Structures. Front Bioeng Biotechnol 2020; 8: 602210. 2020/12/18. DOI: 10.3389/fbioe.2020.602210.

24. Morin KT and Tranquillo RT. In vitro models of angiogenesis and vasculogenesis in fibrin gel. Exp Cell Res 2013; 319: 2409–2417. 2013/06/27. DOI: 10.1016/j.yexcr.2013.06.006.

25. Schwanzer-Pfeiffer D, Rossmanith E, Schildberger A, et al. Characterization of SVEP1, KIAA, and SRPX2 in an in vitro cell culture model of endotoxemia. Cell Immunol 2010; 263: 65–70. 2010/03/20. DOI: 10.1016/j.cellimm.2010.02.017.

26. Nagy JA, Dvorak AM and Dvorak HF. Vascular hyperpermeability, angiogenesis, and stroma generation. Cold Spring Harb Perspect Med 2012; 2: a006544. 2012/02/23. DOI: 10.1101/cshperspect.a006544.

27. Sang Y, Roest M, de Laat B, et al. Interplay between platelets and coagulation. Blood Rev 2021; 46: 100733. 2020/07/20. DOI: 10.1016/j.blre.2020.100733.

28. Hubbard B, Buczek-Thomas JA, Nugent MA, et al. Fibronectin Fiber Extension Decreases Cell Spreading and Migration. J Cell Physiol 2016; 231: 1728–1736. 2015/12/02. DOI: 10.1002/jcp.25271.

29. Carraher CL and Schwarzbauer JE. Regulation of matrix assembly through rigidity-dependent fibronectin conformational changes. J Biol Chem 2013; 288: 14805–14814. 2013/04/17. DOI: 10.1074/jbc.M112.435271.

30. Senzel L, Gnatenko DV and Bahou WF. The platelet proteome. Curr Opin Hematol 2009; 16: 329–333. 2009/06/25. DOI: 10.1097/MOH.0b013e32832e9dc6.

31. Zhou Y, Chen H, Li H, et al. 3D culture increases pluripotent gene expression in mesenchymal stem cells through relaxation of cytoskeleton tension. J Cell Mol Med 2017; 21: 1073–1084. 2017/03/10. DOI: 10.1111/jcmm.12946.

32. Novoseletskaya E, Grigorieva O, Nimiritsky P, et al. Mesenchymal Stromal Cell-Produced Components of Extracellular Matrix Potentiate Multipotent Stem Cell Response to Differentiation Stimuli. Front Cell Dev Biol 2020; 8: 555378. 2020/10/20. DOI: 10.3389/fcell.2020.555378.

33. Zeiger AS, Loe FC, Li R, et al. Macromolecular crowding directs extracellular matrix organization and mesenchymal stem cell behavior. PLoS One 2012; 7: e37904. 2012/06/01. DOI: 10.1371/journal.pone.0037904.

34. Semon JA, Nagy LH, Llamas CB, et al. Integrin expression and integrin-mediated adhesion in vitro of human multipotent stromal cells (MSCs) to endothelial cells from various blood vessels. Cell Tissue Res 2010; 341: 147–158. 2010/06/22. DOI: 10.1007/s00441-010-0994-4.

35. Silva GV, Litovsky S, Assad JA, et al. Mesenchymal stem cells differentiate into an endothelial phenotype, enhance vascular density, and improve heart function in a canine chronic ischemia model. Circulation 2005; 111: 150–156. 2005/01/12. DOI: 10.1161/01.CIR.0000151812.86142.45.

36. Pill K, Melke J, Muhleder S, et al. Microvascular Networks From Endothelial Cells and Mesenchymal Stromal Cells From Adipose Tissue and Bone Marrow: A Comparison. Front Bioeng Biotechnol 2018; 6: 156. 2018/11/10. DOI: 10.3389/fbioe.2018.00156.

37. Banimohamad-Shotorbani B, Kahroba H, Sadeghzadeh H, et al. DNA damage repair response in mesenchymal stromal cells: From cellular senescence and aging to apoptosis and differentiation ability. Ageing Res Rev 2020; 62: 101125. 2020/07/20. DOI: 10.1016/j.arr.2020.101125.

38. Ruster B, Gottig S, Ludwig RJ, et al. Mesenchymal stem cells display coordinated rolling and adhesion behavior on endothelial cells. Blood 2006; 108: 3938–3944. 2006/08/10. DOI: 10.1182/blood-2006-05-025098.

39. Koutsiaris AG, Tachmitzi SV, Batis N, et al. Volume flow and wall shear stress quantification in the human conjunctival capillaries and post-capillary venules in vivo. Biorheology 2007; 44: 375–386. 2008/04/11.

