## Supplementary figures and images for "Mesenchymal stem cells support human vascular endothelial cells to form vascular sprouts in human platelet lysate-based matrices"

### Supplemental Figure 1

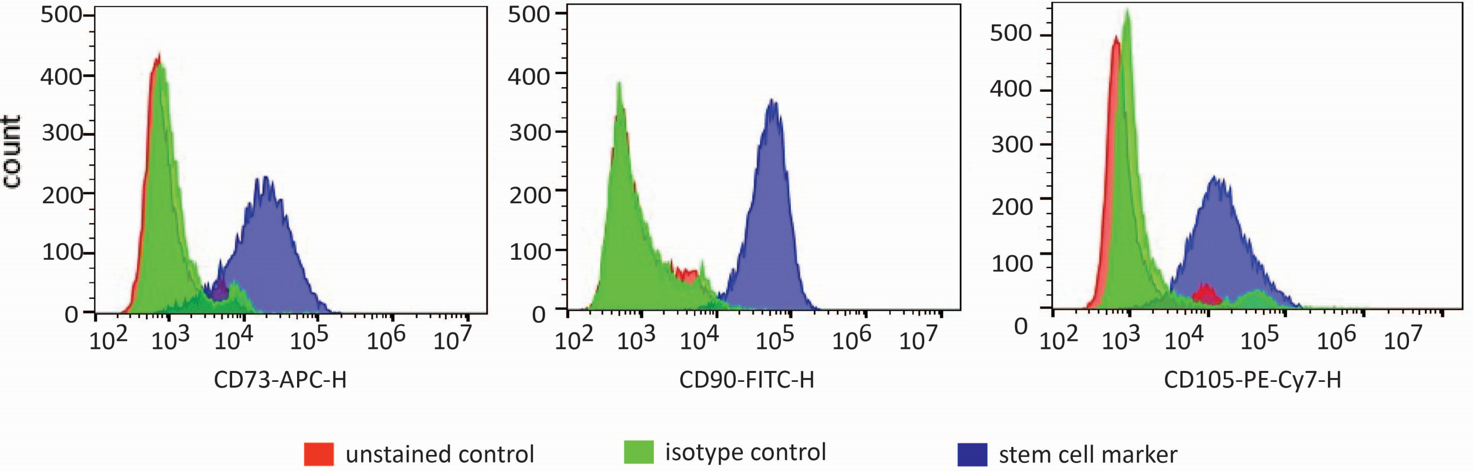

### Supplemental Figure 2

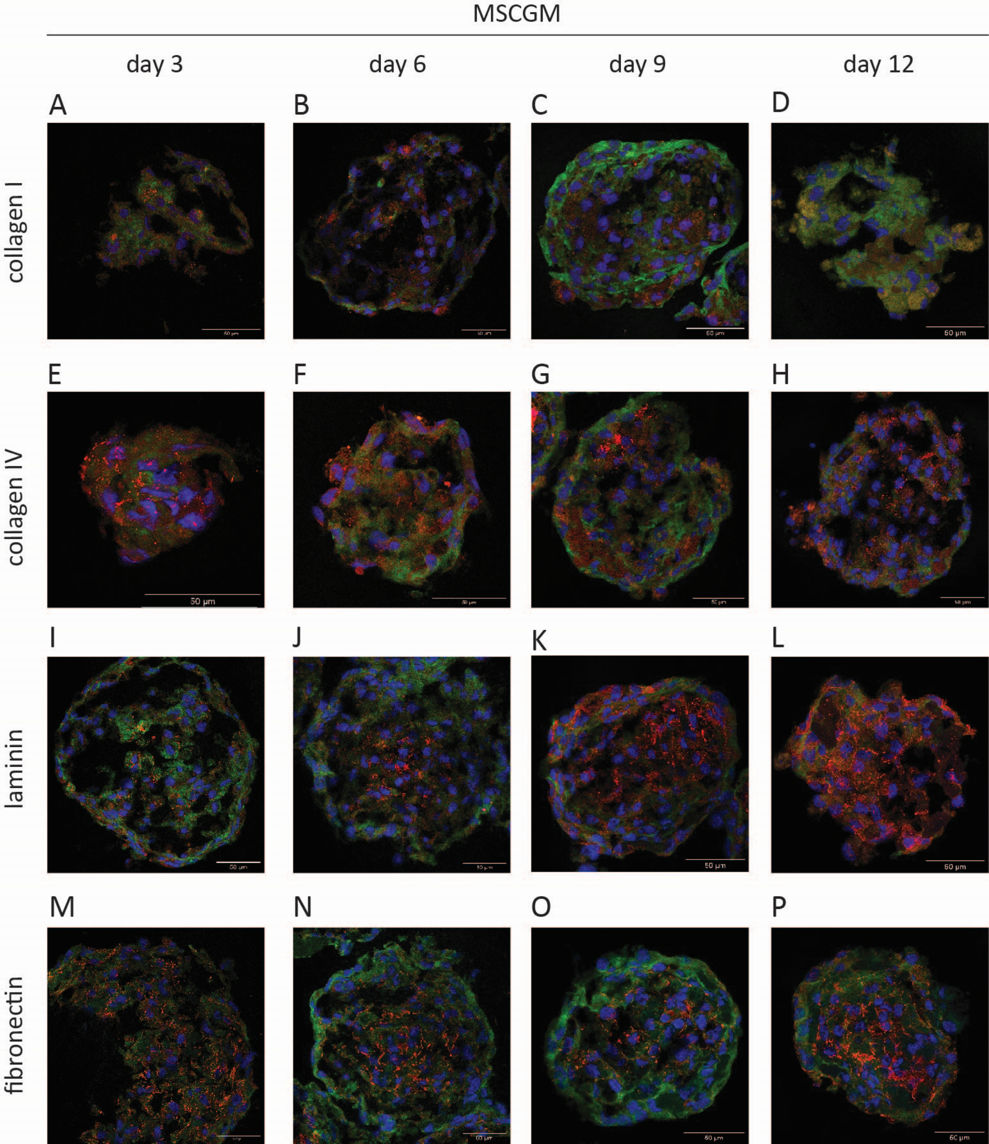

### Supplemental Figure 3

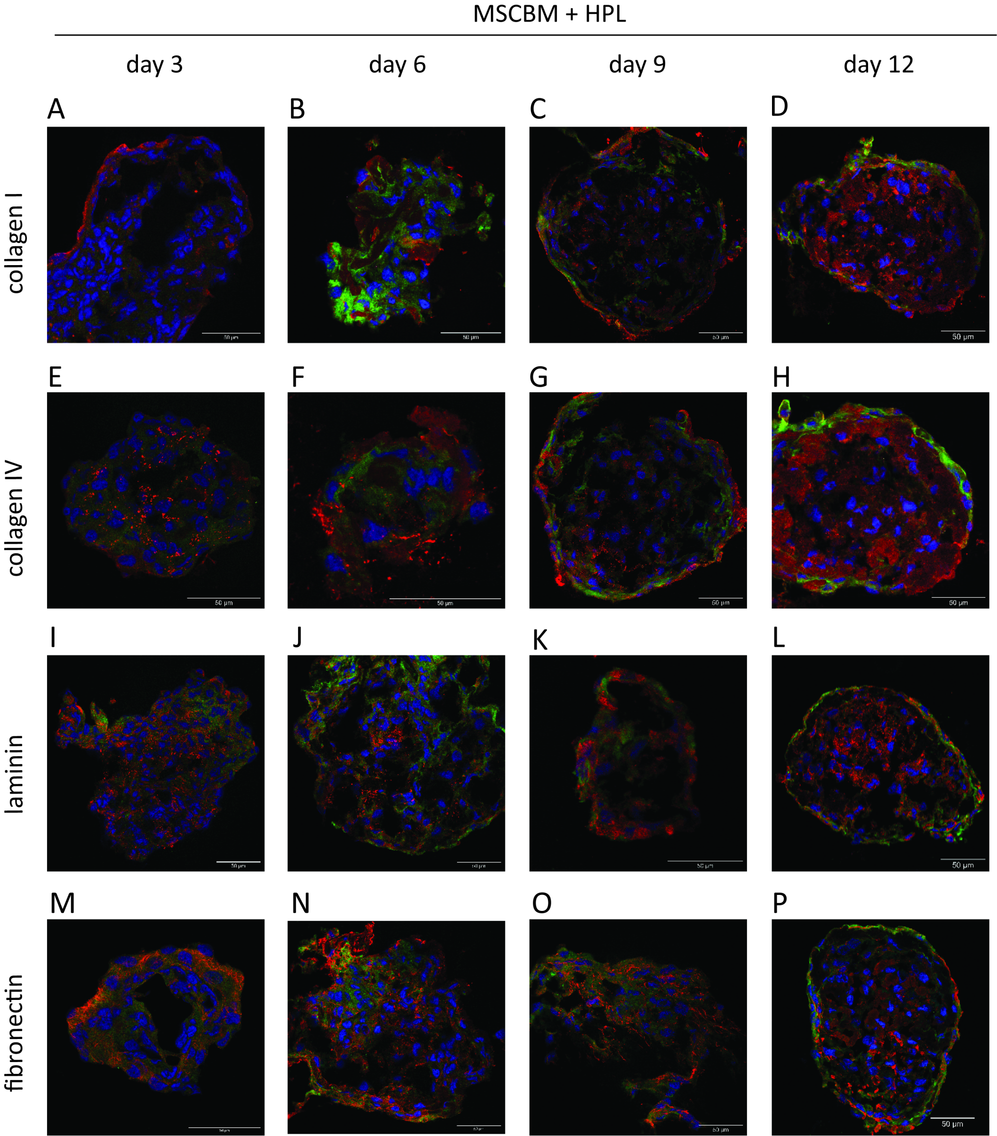
